# An operon encoding two secreted nucleases mediates virulence in Methicillin-resistant *Staphylococcus aureus*

**DOI:** 10.64898/2026.08.26.747337

**Authors:** Alex Zilinskas, Heyuan Michael Ni, Zoe Netter, Kuei-Ho Chen, Danielle L. Swaney, Amir Balakhmet, Nevan Krogan, Sarah Stanley

## Abstract

Methicillin-resistant *Staphylococcus aureus* (MRSA) is an opportunistic pathogen that colonizes a significant proportion of humans, contains numerous virulence factors promoting infection, and continues to threaten human lives and burden healthcare systems globally. Many MRSA virulence factors are known to be either secreted or anchored on the outer leaflet of the cell surface. Although many virulence factors have been studied intensively in MRSA, there remains a significant proportion of secreted and surface proteins that are unstudied for their potential as virulence factors. We began with identifying proteins secreted from MRSA in axenic culture using an unbiased mass-spectrometry based approach. 2 secreted proteins thus identified mapped to an operon of 6 genes, *SAUSA300_1739* to *SAUSA300_1744*. Mutation of each of the individual genes in the operon resulted in attenuation in a mouse model of subcutaneous infection. We demonstrate that two genes in the operon, *SAUSA300_1739*, and *SAUSA300_1740,* encode nucleases with DNase activity. Genetic analysis of the *SAUSA300_1739* to *SAUSA300_1744* operon across several *Staphylococcus aureus* strains indicate that the operon is highly conserved, highlighting its importance for virulence.

**Summary:** Methicillin-resistant *Staphylococcus aureus* (MRSA) remains a serious human pathogen with a myriad of virulence factors supporting pathogenicity. From a MRSA exoproteome, we identified an operon of 6 genes with unknown function that are dispensable for growth in axenic culture but promote MRSA virulence during subcutaneous infection.

## Introduction

*Staphylococcus aureus* is an opportunistic pathogen which colonizes approximately 30-50% of healthy humans (1, 2). *S. aureus* belongs to the “ESKAPE” group of pathogenic bacteria known for their ability to spread in modern hospital care settings and increases in antimicrobial resistant infections (3). In 2017, the more pathogenic and antimicrobial resistant version of *S. aureus*, Methicillin-resistant *Staphylococcus aureus* (MRSA), caused approximately 300,000 hospitalizations, 10,000 deaths, and $1.7 billion in healthcare costs in the United States alone (4). In 2019, MRSA was estimated to cause up to 120,000 deaths worldwide (3). Although MRSA infections can manifest in different tissues, skin and soft tissue infections (SSTIs) are most common, with incidence rates of 32.1 to 48.1 per 1,000 (5). The dominant MRSA clonal type in North America is USA300, which accounts for a large proportion of community and hospital-acquired infections (6).

The global success of *S. aureus* in causing high morbidity and mortality is largely attributable to its arsenal of virulence factors. These virulence factors promote immune evasion, induce host cell lysis, induce non-specific T cell receptor activation, prevent antibody and complement-mediated antimicrobial functions, degrade antibacterial compounds including reactive oxygen species, and inhibit chemotaxis (7). One virulence strategy used by *S. aureus* is the deployment of secreted nucleases. *S. aureus* is known to secrete the DNA nuclease Nuc1 and express a surface-bound nuclease, Nuc2 DNA (8), as well as secrete the EssD DNA nuclease (9) which together contribute to biofilm formation and degrade neutrophil extracellular traps (NETs) to promote infection.

Despite the characterization of many virulence factors, a large percentage of genes in the *S. aureus* pangenome remain unannotated or limited to predicted function. With many genes with unknown or with poorly annotated functions, our inventory of *S. aureus* virulence factors remains incomplete. Uncovering these hidden factors and understanding their impacts on the host is necessary to promote novel vaccine and therapeutic development. Because a large proportion of *S. aureus* virulence factors are secreted or surface-anchored, we focused on characterizing a secreted proteome of the North America-dominant MRSA USA300 clonal type in defined media, including under acidic conditions and hydrogen peroxide stress. We identified a total of 109 secreted proteins, 29 of which have unknown protein function and have not been implicated in *S. aureus* virulence. Using this dataset, we identified 21 transposon mutants, available in a non-essential gene arrayed transposon library, with loss of function mutations in genes without established function and assayed them for virulence defects. We found that two genes, *SAUSA300_1739* and *SAUSA300_1740* are dispensable for growth in liquid culture, however are required for full virulence in a murine subcutaneous infection model. Furthermore, we demonstrate that both SAUSA300*_*1739 and SAUSA300_1740 are nucleases with DNase activity. Finally, we show that every gene in the six gene operon encoding SAUSA300*_*1739 and SAUSA300_1740 contributes to the virulence of MRSA. We find that this operon is highly conserved across MRSA strains, underscoring its importance for virulence.

## Results

### Characterization of MRSA USA300 secreted proteome

We first developed an exoproteome of MRSA USA300 to identify surface and secreted proteins in an unbiased manner. To establish the exoproteome, an overnight culture of WT MRSA USA300 SF8300 strain in minimal media at pH 7.4 was backdiluted to OD_600_ = 0.05 in fresh minimal media at pH 7.4, pH 5.5, or pH 7.4 + 5 mM H_2_O_2_. We chose pH 5.5 as the acidic condition because MRSA USA300 has been shown to prevent phagolysosome maturation with a final pH of ∼5 within the phagosomes of THP-1 macrophages infected with MRSA USA300 (10). Likewise, the 5 mM H_2_O_2_ concentration was chosen as the oxidative stress condition to mimic an oxidative environment of a phagosome. After 1 and 2 hours shaking at 37°C, the supernatants and pellets were processed and characterized via electrospray ionization mass spectrometry. A total of 109 proteins were found to be significantly more abundant in the collected supernatant than the pellet (Supplementary Table 1). Of the 109 proteins, 52 proteins have Sec/SPI signal peptides, 23 proteins have Sec/SPII signal peptides designated for lipoproteins, 6 proteins have been published as secreted proteins, and 9 proteins are transmembrane proteins. 70 proteins have been previously published to have a role as *S. aureus* virulence factors; however, 29 proteins have not been characterized and unknown if they contribute to *S. aureus* virulence during in vivo infection.

### *SAUSA300_1739* and *SAUSA300_1740* have differences in virulence compared to WT MRSA in vitro

Of the 109 proteins collected in this exoproteome, 29 are uncharacterized. Corresponding transposon mutants in the genes of 21 of these proteins were selected from the Nebraska Transposon Mutant Library (NTML) which contains individual arrayed transposon-inserted mutants of all non-essential genes of the *Staphylococcus aureus* genome (11). We first assayed each transposon mutant for the ability to survive during macrophage infection, and for the ability to kill host macrophages. For bacterial CFU determination C57BL/6 (B6) bone-marrow derived macrophages (BMDMs) were infected at MOI of 10 after which gentamicin was added to the media to eliminate extracellular bacteria. For cell killing assays, BMDMs were infected, cultured without gentamicin, and a lactate dehydrogenase (LDH) release assay was used to quantify cell death. Most mutants had no phenotype in either assay (data not shown). Six mutants with transposon insertions in the genes *SAUSA300_1739, SAUSA300_1740, SAUSA300_0883, SAUSA300_2355, SAUSA300_0651,* and *SAUSA300_2315* exhibited differences in CFU and or LDH release assay relative to the WT strains (Supplementary Figure 1 and Figures 1A-D), however these differences were modest.

**Figure 1.**
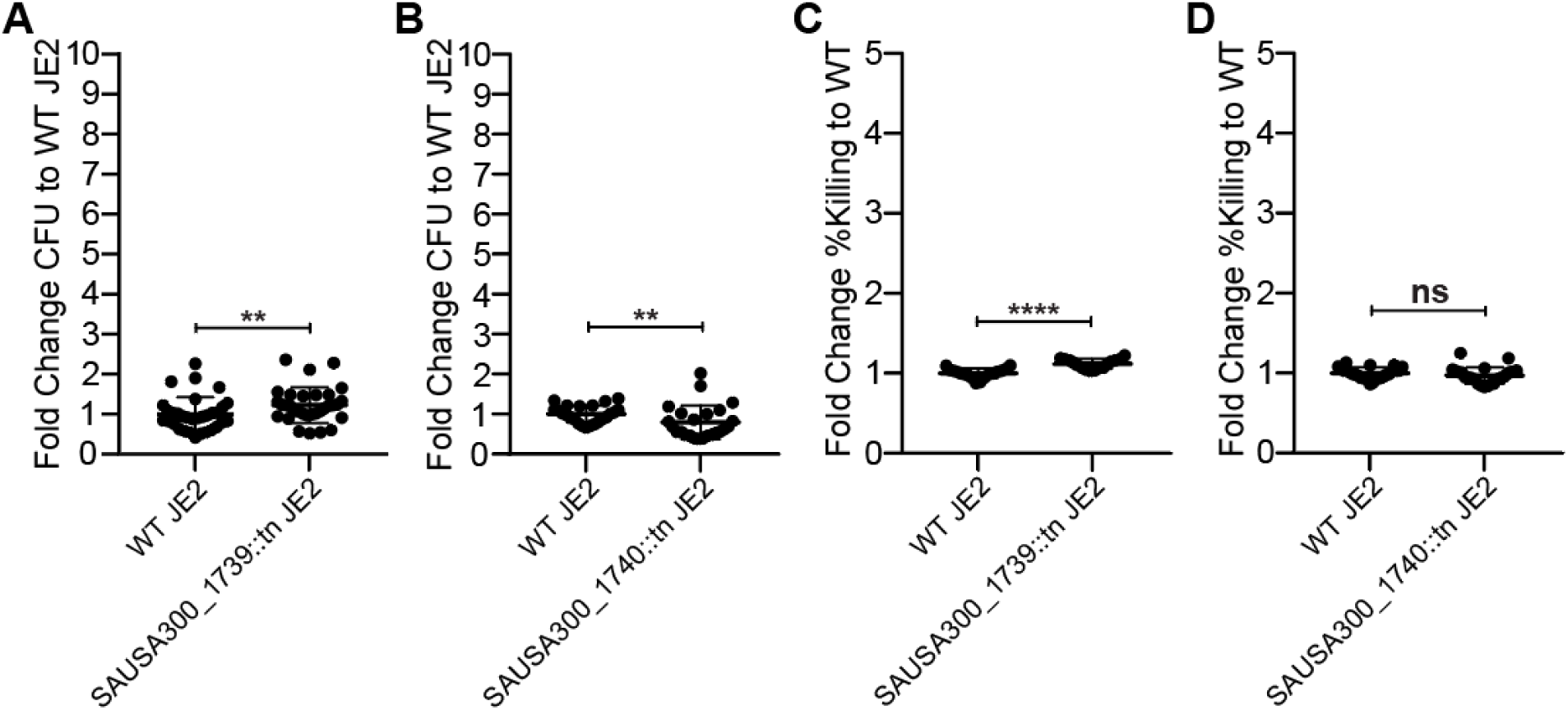
Transposon-inserts of *SAUSA300_1739* and *SAUSA300_1740* show small differences in virulency compared to WT MRSA in vitro. WT, *SAUSA300_1739*::tn, and *SAUSA300_1740*::tn MRSA USA300 JE2 were spinfected into WT C57BL/6 bone-marrow-derived macrophages (BMDMs) at a MOI of 10 followed by gentamicin protection for intracellular MRSA CFU, or BMDMs were infected without spinfection with MRSA at MOI of 10 for Lactate Dehydrogenase (LDH) release assay. **(A, C)** Intracellular CFU of MRSA after 20 hours post spinfection. Fold change CFU results in A and C are from comparison of individual biological replicates against the average WT MRSA USA300 JE2 CFU within the same experiment. **(B, D)** LDH release assay results of MRSA after 6 hours post infection. Fold change %Killing LDH results in B are from comparison of individual biological replicates against the average WT MRSA USA300 JE2 %Killing LDH within the same experiment. **(A)** *SAUSA300_1739* results are from fold change CFU results of 5 pooled experiments and **(B)** *SAUSA300_1740* results are from fold change CFU results of 3 pooled experiments. **(C)** *SAUSA300_1739* results are from fold change LDH results of 2 pooled experiments and **(D)** *SAUSA300_1740* results are from fold change LDH results of 3 pooled experiments. **, p < 0.01; ****, p < 0.0001 (unpaired nonparametric Mann-Whitney U test).

MRSA encodes at least 3 secreted nucleases, Nuc1, Nuc2 (8), and EssD (9) (also called EsaD) that are important for virulence. Nuc1 is secreted, while Nuc2 remains cell associated (8). EssD is secreted through the type VIIb secretion system of *S. aureus* (9). Interestingly, two of the secreted proteins identified in our exoproteome, SAUSA300_1739 and SAUSA300_1740 had protein BLAST search results with some homology to annotated DNA binding proteins and potential nucleases (Supplementary Table 2). These genes are contained within the same predicted operon. We observed statistically significant, but very minor differences in survival in macrophages comparing WT (JE2) with *SAUSA300_1739*::tn and *SAUSA300_1740*::tn mutants (Figure 1A, 1B). Similarly, minimal differences in macrophage cell death were elicited by these mutants relative to WT (Figure 1C, 1D).

### *SAUSA300_1739* and *SAUSA300_1740* belong to an operon of 6 genes with no impact on axenic culture growth rates

Because genes located within an operon are transcriptionally controlled by the same promoter, a reasonable prediction about genes within an operon is that they might be involved in the same cellular processes (12). Since *SAUSA300_1739* and *SAUSA300_1740* are located next to each other in the MRSA USA300 FPR3757 genome, we tested whether *SAUSA300_1739* and *SAUSA300_1740* are transcribed in the same operon. Prokaryote Promoter Prediction v2.0 software predicted that there is only 1 promoter sequence between *SAUSA300_1738* and *SAUSA300_1745*, located upstream of *SAUSA300_1739*, suggesting that the genes between *SAUSA300_1739* to *SAUSA300_1744* may be transcribed together (13) (Figure 2A). To test this, we used reverse transcriptase-PCR (14) of purified WT JE2 MRSA mRNA using primer sets spanning putative gene junctions across the operon (Figure 2A) and found that that *SAUSA300_1739* to *SAUSA300_1744* are encoded on the same mRNA (Figure 2B).

**Figure 2.**
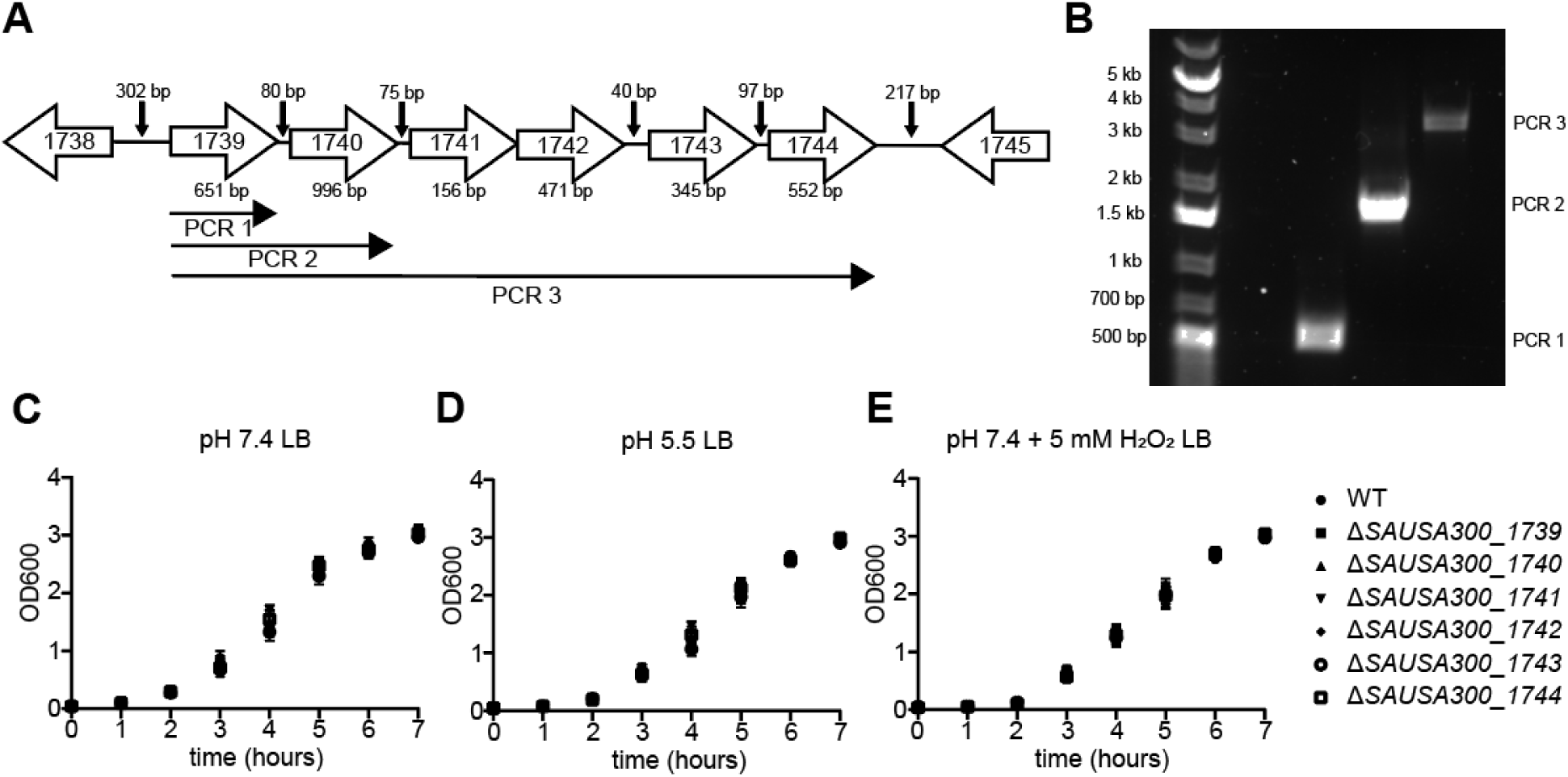
*SAUSA300_1739* and *SAUSA300_1740* are in an operon of 6 genes. WT MRSA USA300 JE2 was grown in LB broth overnight, then backdiluted to OD_600_ = 0.05 and shaken at 37 °C, 180 rpm for 2.5 hours followed by whole RNA extraction and reverse-transcription PCR. **(A)** Representation of gene orientation and distances between *SAUSA300_1738* and *SAUSA300_1745*. **(B)** Agarose gel of reverse-transcription PCR after RNA extraction. Primers for reverse transcription were specific for *SAUSA300_1739*, *SAUSA300_1740*, or *SAUSA300_1744*. PCR of reverse transcription product was performed using forward primer specific for *SAUSA300_1739* and reverse primers specific for either *SAUSA300_1739*, *SAUSA300_1740*, or *SAUSA300_1744*. Results in B represent 3 biological replicates. WT, Δ*SAUSA300_1739*, Δ*SAUSA300_1740*, Δ*SAUSA300_1741*, Δ*SAUSA300_1742*, Δ*SAUSA300_1743*, and Δ*SAUSA300_1744* MRSA USA300 JE2 overnight cultures grown in LB pH 7.4 media was backdiluted into LB media at **(C)** pH 7.4, **(D)** pH 5.5, or **(E)** pH 7.4 with 5 mM H_2_O_2_. OD_600_ was measured every hour for 7 hours. Results in C, D, and E represent triplicate replicates for each timepoint and media conditions.

Thus, *SAUSA300_1739* and *SAUSA300_1740* belong to an operon of six genes from *SAUSA300_1739* to *SAUSA300_1744*. We next generated mutants lacking each individual gene of *SAUSA300_1739* to *SAUSA300_1744* in JE2 using allelic replacement with the pIMAY plasmid (15). All six mutants grew normally in LB broth at pH 7.4, pH 5.5, and pH 7.4+5 mM H_2_O_2_ conditions (Figure 2C, 2D, 2E). These results indicate that loss of any of the 6 genes in the *SAUSA300_1739* to *SAUSA300_1744* operon does not impact growth rate in broth at physiologically relevant pH, acidic, or oxidative stress conditions.

### *SAUSA300_1739* to *SAUSA300_1744* knockouts are attenuated in a mouse subcutaneous infection model

Due to minor differences in MRSA survival within or killing of macrophages in an in vitro model, we investigated if each of these 6 genes has a role in *S. aureus* pathogenesis in vivo, we infected WT B6 mice with WT, mutant, or complemented (generated by transforming each individual mutant with modified expression plasmid pRN11 (16) that had the mCherry reading frame replaced with each of the individual genes PCR amplified from WT MRSA USA300 JE2 chromosomal DNA) strains using a mouse subcutaneous infection model. WT C57BL/6 mice had right lower flanks shaved and were injected with 1 x 10^7^ CFU MRSA. The infection was allowed to proceed for 3 days, at which time lesions were excised using a 12 mm biopsy punch, homogenized, and CFU were enumerated by plating on agar. Surprisingly, each of the mutants had lower lesion CFU of >1 log compared to WT JE2, 3 days post infection (Figure 3A, 3B).

**Figure 3.**
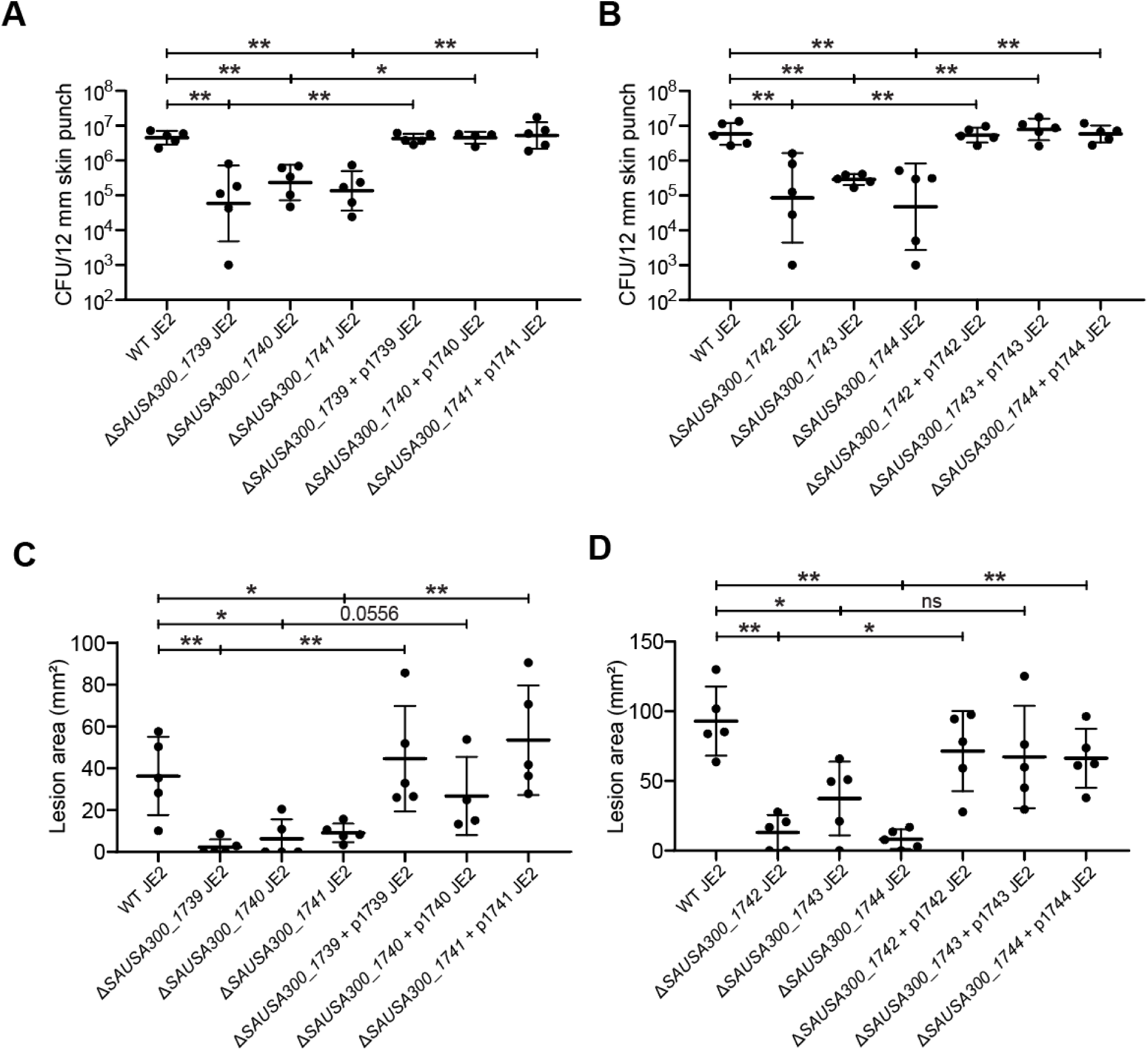
Genetic knockouts of either *SAUSA300_1739* to *SAUSA300_1744* results in attenuation in a murine subcutaneous infection model. WT C57BL/6 mice were infected with 1 x 10^7^ CFU of either WT, Δ*SAUSA300_1739*, Δ*SAUSA300_1740*, Δ*SAUSA300_1741*, complemented Δ*SAUSA300_1739* (+ p1739), complemented Δ*SAUSA300_1740* (+ p1740), or complemented Δ*SAUSA300_1741* (+ p1741) MRSA USA300 JE2, or mice were infected with 1 x 10^7^ CFU of either WT, Δ*SAUSA300_1742*, Δ*SAUSA300_1743*, Δ*SAUSA300_1744*, complemented Δ*SAUSA300_1742* (+ p1742), complemented Δ*SAUSA300_1743* (+ p1743), or complemented Δ*SAUSA300_1744* (+ p1744) MRSA USA300 JE2. **(A, B)** 3 days post infection, CFU from standardized 12 mm skin biopsy punches. **(C, D)** 3 days post infection, open lesion area using caliper. Results in A-D are representative of 2 independent experiments. *, p < 0.05; **, p < 0.01 (unpaired nonparametric Mann-Whitney U test).

Additionally, each of the knockouts caused infections with reduced lesion size area compared to WT and complemented strains at 3 days post infection (Figures 3C, 3D). Taken together, these results indicate that each of the 6 genes in the *SAUSA300_1739* to *SAUSA300_1744* operon is important for *S. aureus* pathogenicity.

### Genetic analysis of SAUSA300_1739 to SAUSA300_1744 across several Staphylococcus aureus strains

The pangenome of *Staphylococcus aureus* is highly diverse with only ∼75% of the genome being conserved between *S. aureus* strains (17). The remaining ∼25% of the *S. aureus* genome consists of mobile genetic elements including pathogenicity islands and plasmids that encode virulence factors (17). Since the ideal candidate for virulence factor-specific therapeutics and vaccine antigens should be highly conserved amongst *S. aureus* strains (18, 19), we examined the conservation of the *SAUSA300_1739* to *SAUSA300_1744* operon among 55 *S. aureus* strains including several community-acquired and hospital-acquired MRSA strains. The *SAUSA300_1739* to *SAUSA300_1744* operon is highly conserved (%nucleotide identity >90%) across the strains. (Figure 4). Interestingly, *SAUSA300_1741* and *SAUSA300_1742*—predicted to encode a lipoprotein and a cytoplasmic protein, respectively—split into two separate genes within the USA300 clonal lineage. Most USA300 (ST8, CC8) strains examined (SUR1, FPR3757, LAC, TCH1516, and JK3137) carry *SAUSA300_1741* and *SAUSA300_1742* as distinct genes, whereas the closely related USA300 strain C8897 (20) retains the ancestral single-gene configuration. This architecture matches what is found in the majority of *S. aureus* strains tested. Additionally, we examined the conservation of the *SAUSA300_1739* to *SAUSA300_1744* operon amongst *Staphylococcus* species. Amongst all the *Staphylococcus* species tested, the *SAUSA300_1739* to *SAUSA300_1744* operon is only found in *S. aureus* and closely related *Staphylococcus* species indicating that the *SAUSA300_1739* to *SAUSA300_1744* operon is evolutionarily new in *Staphylococcus* (Supplementary Figure 2). These results indicate that the *SAUSA300_1739* to *SAUSA300_1744* operon is generally conserved across *S. aureus* strains, consistent with an important role in virulence.

**Figure 4.**
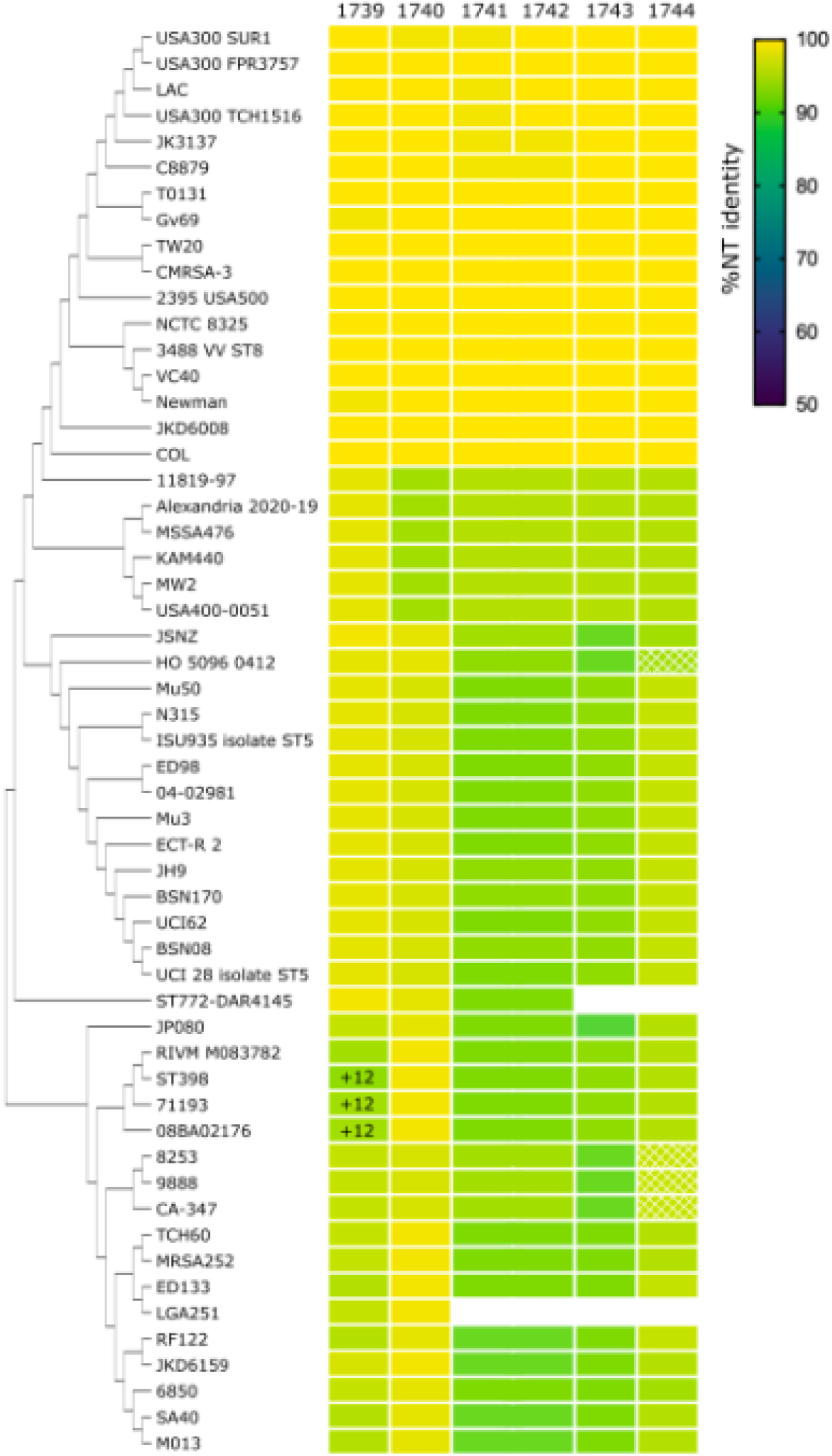
Genetic analysis of *SAUSA300_1739* to *SAUSA300_1744* operon. Whole genome sequences were obtained through the NIH National Center of Biotechnology Information (NCBI) repository, accession numbers listed in materials and methods section. Lack of box for gene indicates lack of gene in strain. “+12” indicates 12 base-pair in-frame insertion in gene of strain. Box with cross hatch marks indicates coding sequence is fully present but interrupted by some insert.

### SAUSA300_1739 and SAUSA300_1740 has DNase activity

SAUSA300_1739 and SAUSA300_1740 are homologous to DNase and DNA-binding proteins (Supplementary Table 2). To test if SAUSA300_1739 and SAUSA300_1740 has DNase activity, we purified both proteins expressed in BL21 *E. coli* under IPTG induction along with Ni-NTA resin to purify 6xHIS-tagged proteins followed by further purification with a FPLC (Supplementary Figure 3). Purified mature unacylated SAUSA300_1739 or SAUSA300_1740 were incubated at 1,000 nM, 2,000 nM, or 5,000 nM of purified with 100 ng of linearized DNA with or without 25 mM EDTA in a Magnesium-containing reaction buffer for indicated times. SAUSA300_1739 had demonstrable DNase activity at 5000nM (Figure 5A) that was abolished in the presence of EDTA, a divalent cation chelator and inhibitor of DNase (21) (Figure 5A).

**Figure 5.**
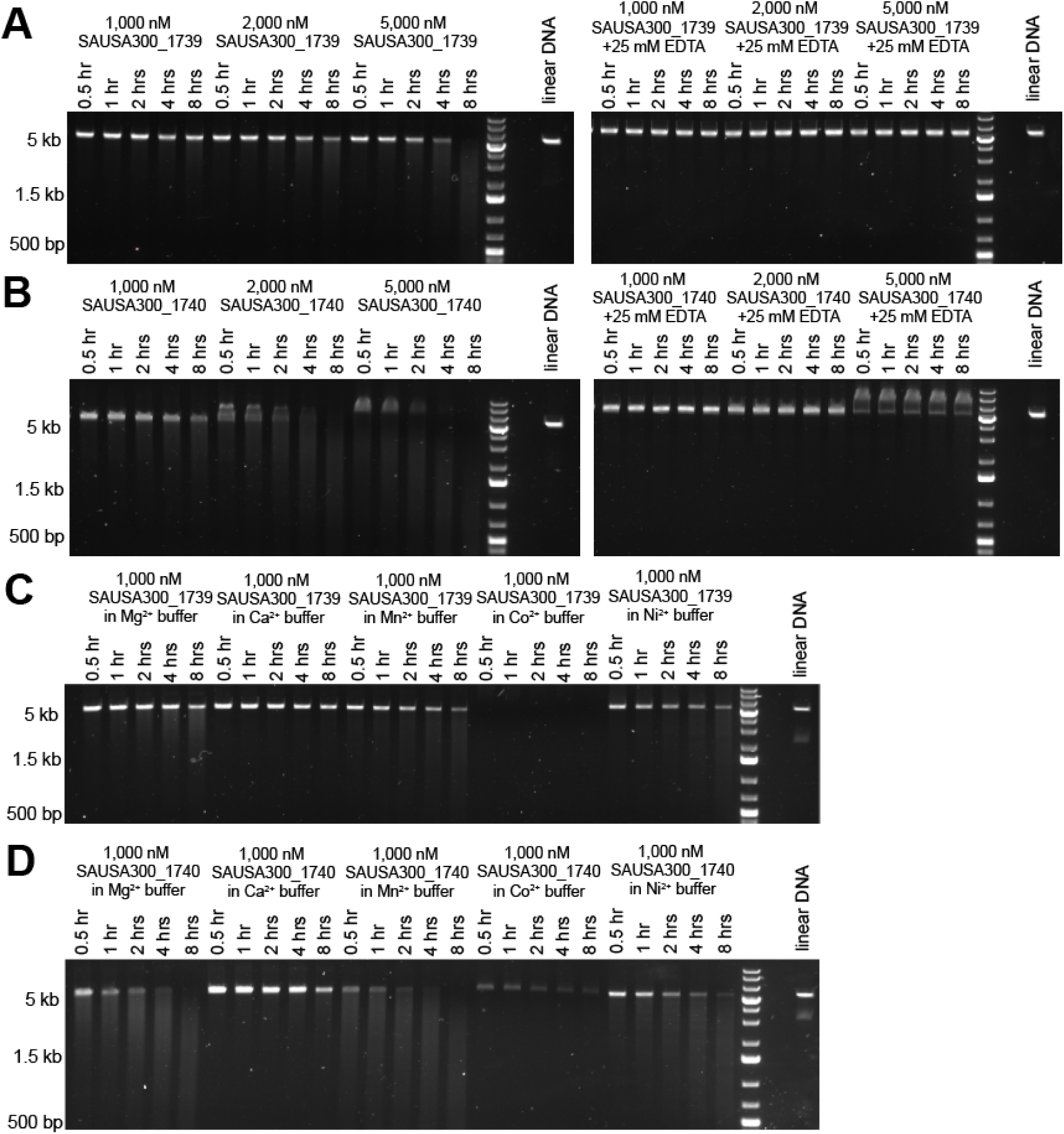
SAUSA300_1739 and SAUSA300_1740 have DNase activity. (A and. **B)** 1,000, 2,000, or 5,000 nM purified mature unacylated SAUSA300_1739 **(A)** or SAUSA300_1740 **(B)** was incubated with 100 ng of linear modified pRN11 plasmid (6 kb size) for indicated times at 37 °C in 3 mM Magnesium-containing reaction buffer without EDTA or with 25 mM EDTA. **(C and D)** 1,000 nM purified unacylated SAUSA300_1739 **(C)** or SAUSA300_1740 **(D)** was incubated with 100 ng of linear modified pRN11 plasmid (6 kb size) for indicated times at 37 °C in 3 mM Mg^2+^, Ca^2+^, Mn^2+^, Co^2+^, or Ni^2+^ containing buffer. Each agarose gel image is representative of 3 independent experiments.

SAUSA300_1740 showed somewhat greater DNase activity with DNAse activity observed at 1000nM (Figure 5B). The addition of EDTA also inhibited DNase activity of SAUSA300_1740 (22) (Figure 5B). Since nucleases can have different levels of nuclease activity based on the cofactor used, we decided to test SAUSA300_1739 and SAUSA300_1740 DNase activity in different cofactor-containing buffers. Interestingly, SASUA300_1739 had DNase activity in Mg^2+^, Mn^2+^, Co^2+^, and Ni^2+^ with no DNase activity observed in Ca^2+^ buffer and a strong DNase activity in Co^2+^ buffer (Figure 5C) with all DNase activity lost when EDTA was present in solution (Supplementary Figure 4). A similar effect was observed with SAUSA300_1740 having DNase activity in Mg^2+^, Ca^2+^, Mn^2+^, Co^2+^, and Ni^2+^ buffer with a weak DNase activity observed in Ca^2+^ buffer and a strong DNase activity in Co^2+^ buffer (Figure 5D), and all DNase activity lost when EDTA was present in solution (Supplementary Figure 4). These results support the idea that SAUSA300_1739 and SAUSA300_1740 are secreted nucleases that promote the virulence of MRSA.

## Discussion

The opportunistic pathogen *Staphylococcus aureus* remains a global burden causing serious and deadly infections. Approximately 20% of humans are persistent carriers, while an additional 60% of humans are intermittently colonized with *S. aureus* reinforcing the importance of studying this major pathogen (23). The rise of antibiotic resistance, the decline of novel antibiotic therapeutic development, and the lack of a protective vaccine all contribute to *S. aureus* remaining a leading cause of human infection. In addition, *S. aureus* is known to utilize an arsenal of virulence factors to evade and manipulate the immune system, lyse host cells, breakdown host antimicrobial molecules, among other mechanisms (7). A greater understanding of how important virulence factors promote *S. aureus* pathogenicity is necessary for anti-virulence factor therapeutic and vaccine development.

To find new virulence factors used by *S. aureus* to promote infection we first focused on identifying uncharacterized proteins secreted by the bacteria. Using this approach, we identified SAUSA300_1739 and SAUSA300_1740 as unknown-function lipoproteins secreted by MRSA JE2. We discovered that *SAUSA300_1739* and *SAUSA300_1740* are members of an operon of 6 genes, *SAUSA300_1739* to *SAUSA300_1744*. The loss of any of these 6 genes does not impact growth of MRSA in axenic culture including in acidic or oxidative conditions (24). Similarly, these genes do not play a large role in the ability of MRSA to survive inside of or kill host cells. Interestingly, *SAUSA300_1739* and *SAUSA300_1740* transposon-inserted MRSA USA300 mutants were hits identified in a screen for mutants that induce less cell damage in an in vitro human endothelial cell line infection model (25), suggesting that these 2 genes have some role as virulence factors. However, there was no additional follow-up on these specific hits in the published study. We observed only minor differences in assays for macrophage killing, suggesting that regulation of cell death may not be a major function of these virulence factors.

Additionally, we show for the first time that mutants lacking *SAUSA300_1739* and *SAUSA300_1740* are attenuated for virulence in vivo. Taken together, the data collectively suggests that these virulence factors likely impact responses that occur in vivo and have less of an impact in cell-based infections in vitro, a finding that is consistent with the hypothesis that these proteins function as nucleases.

In addition to *SAUSA300_1739* and *SAUSA300_1740*, we found that every gene in the operon is required for full virulence of MRSA in skin infections. Previously, a transposon-insertion mutant of *SAUSA300_1741* MRSA USA300 was reported to be attenuated in a murine subcutaneous infection model while not impacting in vitro growth (26). However, to our knowledge, nothing is known about the other genes in the operon, and their possible roles in virulence. Importantly, we found that the *SAUSA300_1739* to *SAUSA300_1744* operon is generally conserved in *S. aureus* strains. *SAUSA300_1740* in particular is highly conserved across majority of analyzed *S. aureus* strains.

*SAUSA300_1740*, a now confirmed expressed lipoprotein virulence factor, could be an ideal candidate for anti-virulence factor therapeutics and antigen for antibody-mediated vaccine due to its presence of the outside of the bacterial cell surface, inhibition of virulence factor mechanism, and conservation in the *S. aureus* genome (17, 18). Future works focused on antibody or small molecule drug-mediated inhibition of SAUSA300_1740 as well the antigenicity of SAUSA300_1740 could improve *S. aureus* infection treatment and vaccine development.

Previously there was almost nothing known about the functions of *SAUSA300_1739* and *SAUSA300_1740.* Protein blast searches suggested both proteins have DNase or DNA-binding activity. Our data indicates that *SAUSA300_1739* and *SAUSA300_1740* both encode DNases. *S. aureus* is known to utilize surface and secreted nucleases to regulate biofilm formation for immune evasion, to breakdown antimicrobial neutrophil extracellular traps (NETs) and to produce cytotoxic concentrations of deoxyribonucleotides (8, 27). In the future, it will be important to identify the redundancies and differences in biochemical mechanisms of *Staphylococcus aureus* secreted nucleases nuc1, nuc2 (8), EssD (9), and newly identified SASUA300_1739, and SAUSA300_1740. In summary, we have identified that the *SAUSA300_1739* to *SAUSA300_1744* operon encodes 6 novel virulence factors with *SAUSA300_1739* and *SAUSA300_1740* having DNase activity and the remaining 4 genes with unknown function. From this work, our understanding of the *S. aureus* virulence factors important for establishing skin and soft tissue infections expands. Moving forward, it will be important to discover the functions of the 4 remaining genes of this operon and figure out the relationship between all 6 genes.

## Materials and Methods

### Ethics statement

All procedures involved with the use of mice were approved by the University of California, Berkeley’s Institutional Animal Care and Use Committee (protocol 2015-09-7979). All protocols conform to federal regulations, the National Research Council Guide for the Care and Use of Laboratory Animals, and the Public Health Service Policy on Humane Care and Use of Laboratory Animals.

### Bacterial culture for mass spectrometry

Minimal media at pH 7.4 was inoculated with WT Methicillin-resistant *Staphylococcus aureus* USA300 strain SF8300 and grown overnight at 37 °C, 180 rpm. Media was prepared with (mg/mL): K₂HPO₄ (70), KH₂PO₄ (20), and (NH₄)₂SO₄ (10), along with the amino acids phenylalanine (4), isoleucine (3), tyrosine (5), cysteine (2), glutamate (10), lysine (1), methionine (7), histidine (3), tryptophan (1), leucine (9), aspartate (9), arginine (7), serine (3), alanine (6), threonine (3), glycine (5), valine (8), and proline (1). It also included the nucleobases adenine (0.5), cytosine (0.5), guanine (0.5), thymine (2), and uracil (0.5). Additional components were MgSO₄·7H₂O (5), FeCl₃ (8), thiamine (1), niacin (1.2), biotin (0.005), calcium pantothenate (0.25), ZnCl₂ (0.07), MnCl₂·4H₂O (0.099), boric acid (0.006), CoCl₂·6H₂O (0.397), CuCl₂·2H₂O (0.003), NiCl₂·6H₂O (0.024), Na₂MoO₄·2H₂O (0.036), and glucose (5).The next day, overnight culture was back-diluted to OD_600_ = 0.05 into fresh minimal media at pH 7.4, pH 5.5, or pH 7.4+5 mM H_2_O_2_. Back-diluted cultures were shaken at 37°C, 180 rpm for 1 to 2 hours before extraction of culture filtrate and cells for mass spectrometry.

### Proteomics sample preparation

Protein was precipitated from 100 μL of each sample (either filtered media or bacterial cell lysate) by the addition of 900 μL of methanol, followed by centrifugation at 21,130 x g for 10 minutes. The supernatant of this precipitation was removed from the tube, leaving only the precipitated protein pellet, which was then resuspended for 5 min at 95 °C with shaking (1,000 rpm) in a 50 μL of 1% sodium deoxycholate in a 100 mM Tris-HCl buffer at pH 8.0. Following protein concentration determination by a Bradford assay, 30 μg of protein from each sample was then transferred to a new tube and reduction and alkylation of cysteine disulfide bonds was performed by the addition of 10 mM TCEP (tris(2-carboxyethyl)phosphine) and 40 mM 2-chloroacetamide for 5 min at 45 °C with shaking (1,500 rpm). Digestion of proteins into peptides was initiated by the addition of 1 μg of LysC (Wako) and 0.5 μg of trypsin (Promega), followed by overnight incubation at 37 °C. Digestion was quenched by the addition of 10% trifluoroacetic acid until a pH of ∼2-3 was reached. The digested lysates were clarified by centrifugation for 3 min at 21,130 x g, and the clarified peptide lysate was desalted on C18 stage tips (Nest Group). The resulting desalted peptides were dried by vacuum centrifugation, resuspended in 0.1% formic acid prior to MS analysis.

### Mass spectrometry data acquisition and analysis

Approximately 0.6μg of sample was injected into an Orbitrap Fusion Lumos Tribrid Mass Spectrometer (Thermo Scientific) operated in positive ion mode using an EASY-nLC 1200 UHPLC (Thermo Scientific). Peptides were separated over a 75-μm internal diameter × 25-cm long picotip column packed with 1.9 μm C18 particles (Dr. Maisch) at a flow rate of 300 nL/min. Buffer A was comprised of 0.1% formic acid, and buffer B was comprised of 80% acetonitrile in 0.1% formic acid. Peptides were eluted by a linear gradient of 1 to 28% B over 25 minutes, followed by a 7 min ramp from 28 to 32% B, and finally a 3 min ramp to 95% buffer B and a subsequent 10min wash at 95% buffer B. MS data was collected over a 45 minutes total acquisition time with MS1 detection in the Orbitrap at 240K resolution in profile mode, 250-1350 m/z scan range, a 50 ms maximum injection time, a 1×10^6^ AGC target, a 30% RF lens setting, a 275 °C transfer tube temperature, and a 2 kV spray voltage. Data dependent MS2 scans were performed in centroid mode using Advanced Peak Determination on charge states from 2-6, with a +/- 10 ppm exclusion for a duration of 40 seconds after one observation.

Selected precursors were isolated with a 0.7 m/z window and fragmented with a 32% normalized HCD energy followed by ion-trap detection in rapid scan mode rate, a 200-1200 m/z range, a 20 ms maximum injection time, and a 3×10^5^ AGC target.

Raw data was searched against the *Staphylococcus aureus* (strain USA300) Uniprot protein database (downloaded July 5, 2020) using Maxquant (version 1.6.12.0) (28). Default search parameters were used, including a minimum peptide length of 7 amino acids, variable modification of acetylation of the protein N-terminus, variable modification of oxidation on methionine, match between runs was disabled, and peptides and protein were filtered to a 1% false discovery rate. MS1 area under the curve intensity values were then analyzed by SAINTexpress (29) was used to confidently discriminate secreted bacterial proteins from contaminating cellular proteins. Proteins with a SAINTexpress BFDR < 0.05 were considered to be high-confidence *Staphylococcus aureus* secreted proteins.

### Bone-marrow macrophage infection assay

Overnight MRSA cultures were prepared by inoculating LB broth with WT or transposon-inserted mutant MRSA USA300 JE2 strain and culturing overnight at 37°C, 180 rpm.

∼16 hours later, overnight cultures were back-diluted in fresh LB broth at OD_600_ = 0.05 and cultured for ∼2 hours to mid-log growth (OD_600_ = ∼0.5). At mid-log growth phase, MRSA cultures were washed twice 1x PBS, pH 7.4. Final resuspension of MRSA pellet was with Bone-marrow-derived macrophage (BMDM) media.

For CFU measurements, BMDMs were spin-fected with MRSA in BMDM media at 1,200 rpm for 5 minutes for a MOI = 10. After spin-fection, BMDMs were washed once with 1x PBS, pH 7.4, then incubated at 37 °C, 5% CO_2_ for 20 hours in BMDM media containing 100 μg/mL gentamicin (Sigma-Aldrich G1397). BMDMs were lysed while suspended in Milli-Q water for 10 minutes at room temperature. Intracellular MRSA was plated on LB agar plates and incubated overnight at 37 °C.

For LDH measurements, BMDMs were suspended in BMDM media containing MRSA at a MOI = 10. Uninfected BMBM wells and empty wells were suspended in BMDM media alone. Then, plates were incubated at 37 °C, 5% CO_2_ for 6 hours. 10x (10% v/v) Triton-X in Milli-Q water was added to positive control well. Media from BMDM plates was mixed 1:1 LDH working reagent (20 μL of 36 mg/mL Lithium L-lactate (Sigma Aldrich L2250) in 10 mM Tris buffer, pH 8.5; 2 μL of 20 mg/mL Iodonitrotetrazolium chloride (Sigma Aldrich I8377) in DMSO; 2 μL of 13.5 U/mL Diaphorase (Roche 10411558001), 3 mg/mL NAD+ (Sigma Aldrich N3014), 0.03% w/v BSA, 1.2% w/v sucrose in 1x PBS, pH 7.4; and 36 μL of 1% w/v BSA in 1x PBS, pH 7.4; per reaction) and incubated at room temperature for 30 minutes covered from light. LDH release assay reaction products were mixed 1:1 with 10% formalin and measured absorbance at 490 nm.

### RNA extraction, reverse transcription, and PCR

WT MRSA USA300 JE2 culture in LB media was grown overnight at 37 °C, 180 rpm. The next morning, the overnight culture was backdiluted into LB media to OD_600_ = 0.05 then shaken at 37 °C, 180 rpm for 2 hours and 30 minutes. MRSA cells were pelleted and lysed with TRIzol reagent (Invitrogen 15596026), bead beaten with 0.1 mm silica beads (MP 116911100) using Mp FastPrep-24 Bead Beater (MP 116004500). Bead beaten TRIzol cell extracts were given chloroform and phase separated with centrifugation at 12,000 rcf, 10 minutes, 4 °C. RNA was extracted and purified using the RNeasy mini kit (Qiagen 74104) with on-column DNase digest. Reverse transcription was performed with Superscript III Reverse Transcriptase kit (Invitrogen 18080093) and reverse transcription primers specific for either SAUSA300_1739, SAUSA300_1740, or SAUSA300_1744 (Table 1). Reverse transcription product was used for PCR reaction with Q5 DNA polymerase kit (NEB M0491L) with forward primer specific for SAUSA300_1739 and reverse primers specific for either SAUSA300_1739, SAUSA300_1740, or SAUSA300_1744. PCR products were run on 1% w/v agarose gel in TAE buffer.

**Table 1.**
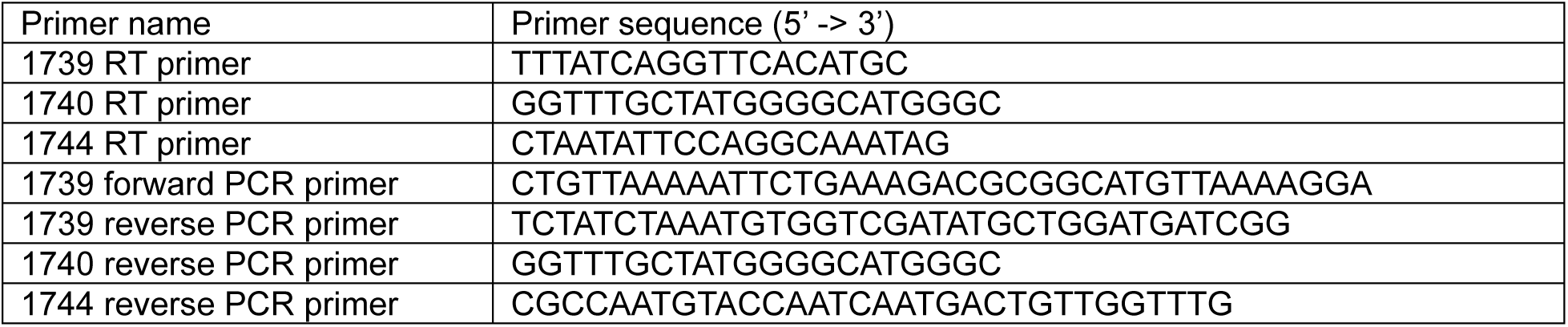
Primer sequences for reverse transcription and PCR.

### Genetic analysis of SAUSA300_1739 to SAUSA300_1744 operon

Phylogenetic tress of *S. aureus* strains were constructed using the tool VBCG (30) with default parameters. *S. aureus* operon *SAUSA300_1739-SAUSA300_1744* nucleotide coding sequences from USA300 FPR3757 strain were used to query an NCBI BLAST database of *S. aureus* reference genomes to identify the operon in other strains.

Heatmaps depicting percent identity were generated in Prism.

Accessed online through the National Center for Biotechnology Information (NCBI) repository whole genome sequences of *Staphylococcus aureus* strains: COL (NCBI accession number: CP000046), JP080 (NCBI accession number: AP017922.1), KAM440 (NCBI accession number: AP040133.1), Newman (NCBI accession number: AP009351.1), MRSA252 (NCBI accession number: BX571856.1), JKD6008 (NCBI accession number: CP002120.1), MSSA476 (NCBI accession number: NC_002953.3), NCTC 8325 (NCBI accession number: NC_007795.1), 04-02981 (NCBI accession number: NC_017340.1), 6850 (NCBI accession number: NC_022222.1), ED133 (NCBI accession number: NC_017337.1), JKD6159 (NCBI accession number: NC_017338.2), Mu3 (NCBI accession number: AP009324.1), RF122 (NCBI accession number: NC_007622.1), TCH60 (NCBI accession number: CP002110.1), VC40 (NCBI accession number: NC_016912.1), N315 (NCBI accession number: NC_002745.2), USA300_FPR3757 (NCBI accession number: CP000255.1), 08BA02176 (NCBI accession number: NC_018608.1), 71193 (NCBI accession number: NC_017673.1), ED98 (NCBI accession number: NC_013450.1), JH9 (NCBI accession number: NC_009487.1), LGA251 (NCBI accession number: FR821779.1), Mu50 (NCBI accession number: NC_002758.2), ST398 (NCBI accession number: NC_017333.1), TW20 (NCBI accession number: FN433596.1), JSNZ (NCBI accession number: CM129921.1), 11819-97 (NCBI accession number: NC_017351.1), EXT-R 2 (NCBI accession number: NC_017343.1), HO 5096 0412 (NCBI accession number: NC_017763.1), M013 (NCBI accession number: NC_016928.2), MW2 (NCBI accession number: NC_003923.1), T0131 (NCBI accession number: NC_017347.1), USA300_TCH1516 (NCBI accession number: NC_010079.1), LAC (NCBI accession number: NZ_CP055225.1), C8879 (NCBI accession number: NZ_CP020956.1), JK3137 (NCBI accession number: NZ_CP020960.1), USA400-0051 (NCBI accession number: CP019574.1), 2395 (NCBI accession number: NZ_CP007499.1), CA-347 (NCBI accession number: NC_021554.1), CMRSA-3 (NCBI accession number: NZ_CP029685.1), UCI 28 isolate ST5 (NCBI accession number: NZ_CP018768.1), UCI62 (NCBI accession number: NZ_CP018766.1), ISU935 isolate ST5 (NCBI accession number: NZ_CP017090.1), SA40 (NCBI accession number: NC_022443.1), USA300_SUR1 (NCBI accession number: CP009423.1), 3488_VV_ST8 (NCBI accession number: NZ_CP089502.1), Gv69 (NCBI accession number: CP009681.1), BSN170 (NCBI accession number: NZ_CP151257.1), BSN08 (NCBI accession number: NZ_CP186027.1), Alexandria_2020-19 (NCBI accession number: CP113244.1), RIVM_M083782 (NCBI accession number: NZ_CP096532.1), 8253 (NCBI accession number: CP166864.1), 9888 (NCBI accession number: CP166863.1), and ST772-MRSA-V strain DAR4145 (NCBI accession number: CP010526.1).

### Generation of genomic knockouts and complementation

WT MRSA USA300 JE2 genomic DNA was purified using NEB Monarch Genomic DNA Purification Kit (NEB T3010S) and Lysostaphin (Millipore Sigma SRE0053). 1 kb upstream and downstream of coding regions of *SAUSA300_1739* and *SAUSA300_1740* were cloned from MRSA genomic DNA and cloned into pIMAY using NEBridge Golden Gate Assembly Kit (BsaI-HF v2) (NEB E1602) with pIMAY-GG-fwd and pIMAY-GG-rev primers listed in table 2. *SAUSA300_1741* to *SAUSA300_1744* coding regions were cloned from MRSA genomic DNA and cloned into pIMAY using NEB Gibson Assembly Cloning kit (NEB E5510S) and pIMAY-GA-fwd and pIMAY-GA-rev primers listed in table 2. pIMAY constructs containing 1 kb upstream and 1 kb downstream of each gene were first transformed into DH5α, then plasmids from DH5α were transformed into IM08B (31). Plasmids purified from IM08B were electroporated in WT MRSA USA300 JE2 strain. Genetic manipulation of WT MRSA USA300 JE2 strain was performed as described in Monk, et al., 2012, to generate knockouts of *SAUSA300_1739* to *SAUSA300_1744*.

**Table 2.**
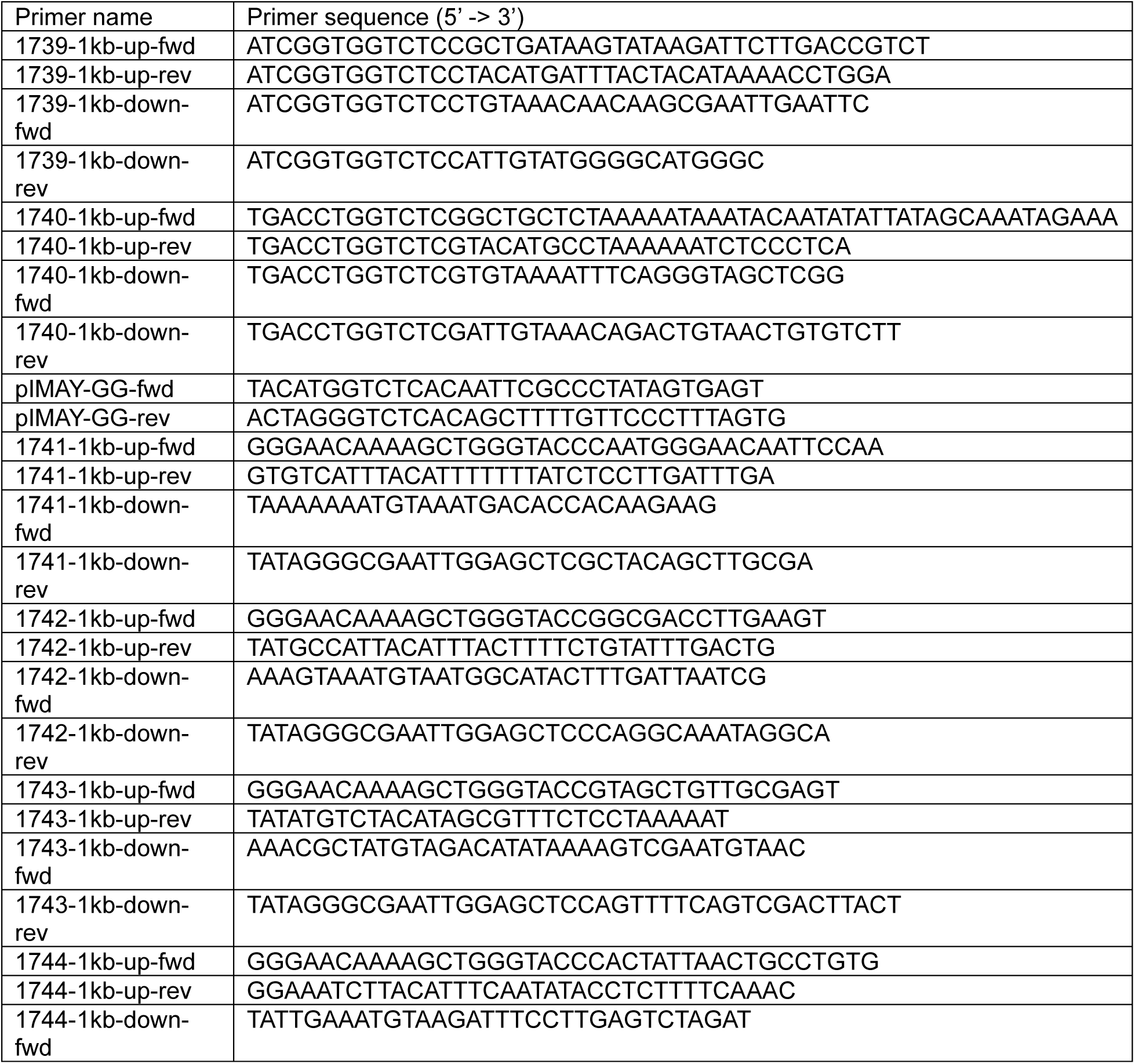
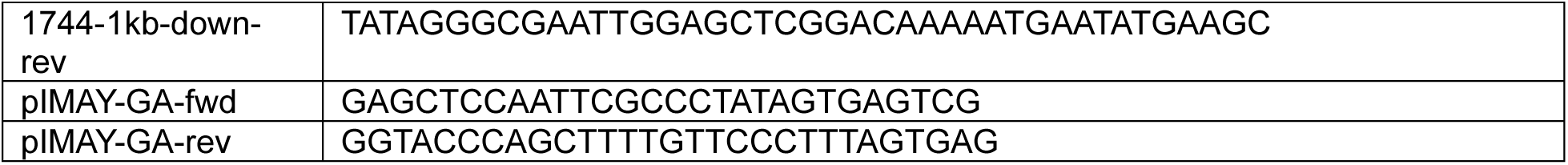
Primer sequences for making knockouts of *SAUSA300_1739* to *SAUSA300_1744*.

Complements were generated by cloning coding regions of each gene *SAUSA300_1739* to *SAUSA300_1744* from purified WT MRSA USA300 JE2 genomic DNA using primers GI-fwd and GI-rev primers, and amplification of pRN11 backbone removing the mCherry gene (16) with primers backbone-fwd and backbone-rev primers listed in Table 3. PCR products were annealed together with NEB Gibson Assembly Cloning kit (NEB E5510S). Plasmids were transformed into DH5α, then plasmids from DH5α were transformed into IM08B (31). Plasmids purified from IM08B were electroporated into genomic knockouts of gene encoded in modified pRN11 expression plasmids (i.e. Δ*SAUSA300_1739* JE2 was electroporated with pRN11-1739).

**Table 3.**
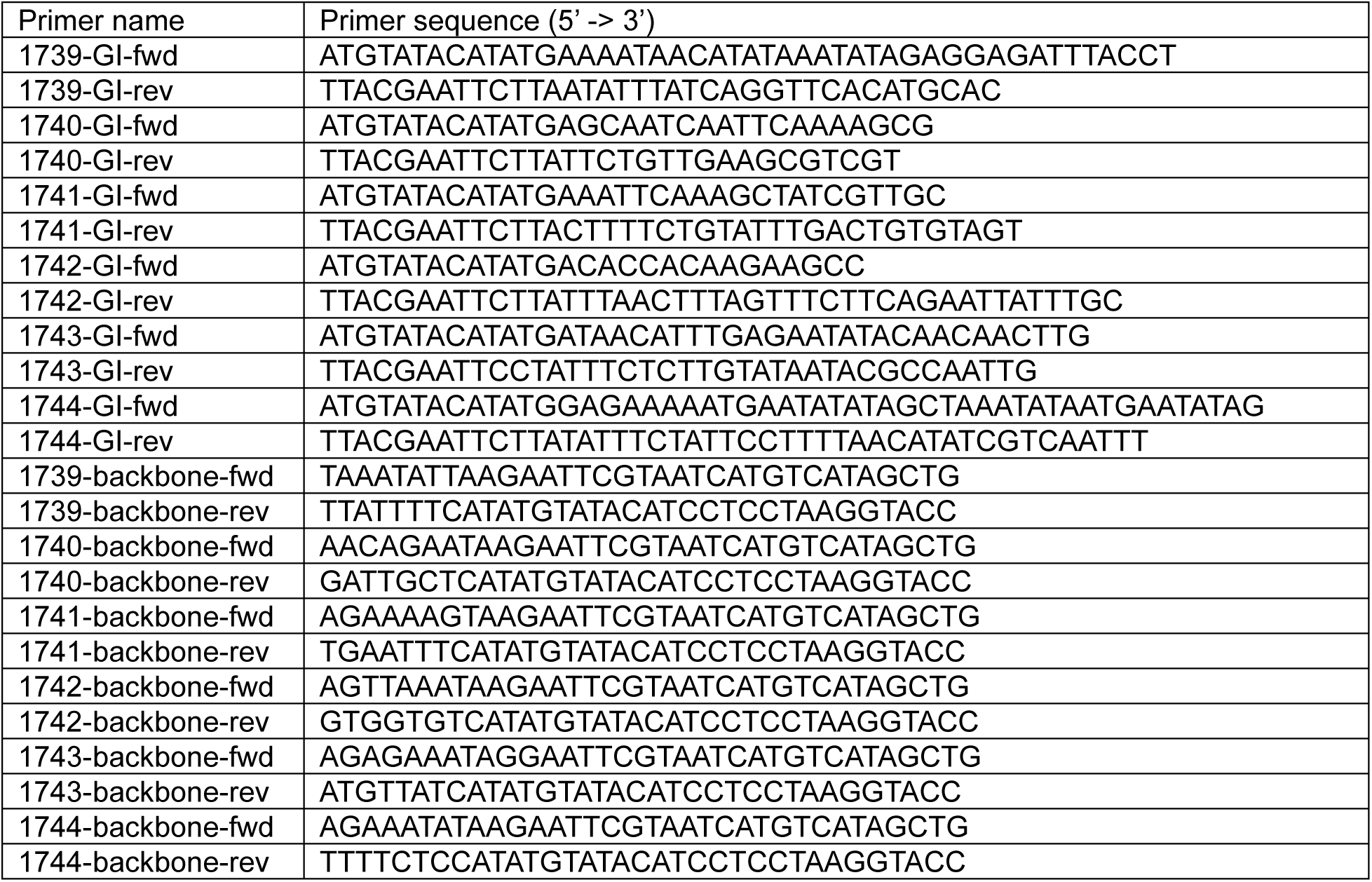
Primer sequences for cloning *SAUSA300_1739* to *SAUSA300_1744* coding regions and amplification of pRN11 backbone for complementation.

### Murine subcutaneous infection model

Day prior to infection, WT C57BL/6 (Inotiv C57BL/6JRccHsd) mice had lower back region between back and right hind leg shaved. Day of infection, overnight MRSA cultures in LB media were back-diluted to OD_600_ = 0.05 in fresh LB media and cultured at 37°C, 180 rpm for 2 hours 15 minutes. Once MRSA culture has reached mid-log growth phase (OD_600_ = ∼0.5), MRSA cells were washed twice with 1x PBS, pH 7.4.

MRSA cells were final resuspension in 1x PBS, pH 7.4 and diluted down to 1 x 10^8^ CFU/mL. Anesthetized mice were subcutaneously injected with 1 x 10^7^ CFU in shaved region of lower right back of mice between spine and right back leg. 3 days post infection, mice were euthanized. Lesion sizes were measured with caliper and extracted with 12 mm biopsy punch (Acuderm P1225). Following bead beating and serial dilutions, homogenized lesion samples were plated on LB agar plates which were incubated overnight at 37 °C.

### Expression of SAUSA300_1739 and SAUSA300_1740 in *E. coli* BL21 cells

WT JE2 MRSA USA300 genomic DNA was PCR’ed to clone *SAUSA300_1739* and *SAUSA300_1740* open-reading frames with removal of the N-terminal sec-Lipo sequence and addition of 6x His-tag on C-terminus. Open-reading frames were cloned into *E. coli* expression plasmid pET14b. pET14b encoding *SAUSA300_1739* or *SAUSA300_1740* were transformed into BL21 cells (NEB C2527H). BL21 colonies with either pET14b-*SAUSA300_1739* or pET14b-*SAUSA300_1740* were inoculated in LB broth with antibiotic selection and grown at 37 °C, 180 rpm. Once cultures reached mid-log growth, IPTG was added at 1 mM final concentration and cultures were shaken at 16 °C, 180 rpm overnight.

After overnight induction, BL21 cells were lysed with CelLytic B cell Lysis Reagent (Sigma-Aldrich B7435). Cell lysate was incubated with 5 mL Ni-NTA resin (GoldBio H-350-50) in 20 mL gravity column. Ni-NTA beads were washed twice with 20 mL first wash buffer (25 mM Tris-HCl, 150 mM NaCl, 20 mM Imidazole, pH 7.6), then washed once with 20 mL second wash buffer (25 mM Tris-HCl, 150 mM NaCl, 40 mM Imidazole, pH 7.6), and eluted with 10 mL elution buffer (25 mM Tris-HCl, 150 mM NaCl, 350 mM Imidazole, pH 7.6). Protein was concentrated and buffer exchanged into final suspension buffer (100 mM NaCl, 25 mM Tris-HCl, 10% v/v glycerol, pH 7.6) using centrifugal concentrator (Millipore UFC9003). Proteins were further purified using a Cytiva Akta Pure 25 FPLC with Superdex 200 Increase 10/300 GL (size exclusion column) running at 0.25 mL/minute at 4 °C in final suspension buffer.

### DNase activity assay

Modified pRN11 plasmid, pAZ14, was linearized using EcoRI digestion (NEB R3101L) to make a 6 kb linear DNA product. 100 ng of linearized pAZ14 was incubated in reaction buffer (3 mM MgCl_2_, 50 mM NaCl, 10 mM Tris-HCl, pH 8.0) with purified SAUSA300_1739 or SAUSA300_1740 at designated concentrations (1,000 nM, 2,000 nM, or 5,000 nM) with or without EDTA (25 mM) and incubated at 37 °C for designated timepoints (0.5 hours, 1 hour, 2 hours, 4 hours, or 8 hours). For other cofactor-containing buffers, 3 mM MgCl_2_ was replaced with either 3 mM CaCl_2_, 3 mM MnCl_2_ x 4H_2_O, 3 mM CoCl_2_ x 6H_2_O, or 3 mM NiCl_2_. DNase reaction products ran on 1% w/v agarose gels in TAE buffer at 100 V for 45 minutes.

## Data availability

All raw MS data files, search results, and individual spectral libraries were uploaded in private mode to the Pride partner ProteomeXchange repository (32) under the identifier PXD072065.

## Acknowledgements

We acknowledge funding from the National Institutes of Health grant U19AI135990 to N.J.K. We thank Binh Diep’s lab for the kind gift of WT MRSA USA300 SF8300 strain. The WT MRSA USA300 JE2 strain was provided by the Network on Antimicrobial Resistance in *Staphylococcus aureus* (NARSA) for distribution through BEI Resources, NIAID, NIH: *Staphylococcus aureus* subsp. *aureus*, Strain JE2, NR-46543. The Nebraska Transposon mutant library was provided by the Network on Antimicrobial Resistance in *Staphylococcus aureus* (NARSA) for distribution through BEI Resources, NIAID, NIH: Nebraska Transposon Mutant Library (NTML) Screening Array, NR-48501. We thank Eva Harris’ lab, especially Elias Duarte and Nharae Lee, for access and their help with FPLC.

## Supplementary Data

**Supplementary Table 1.**
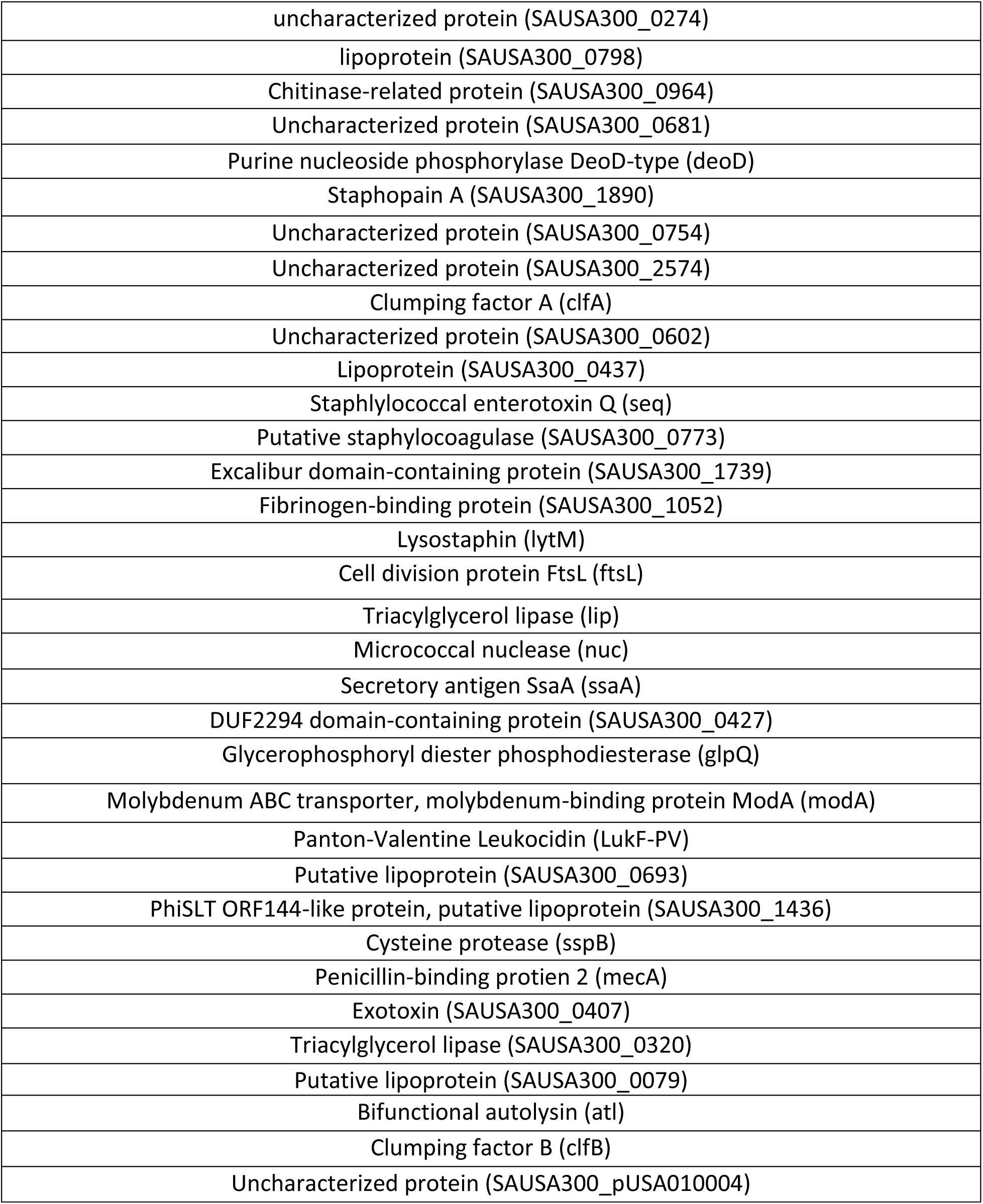
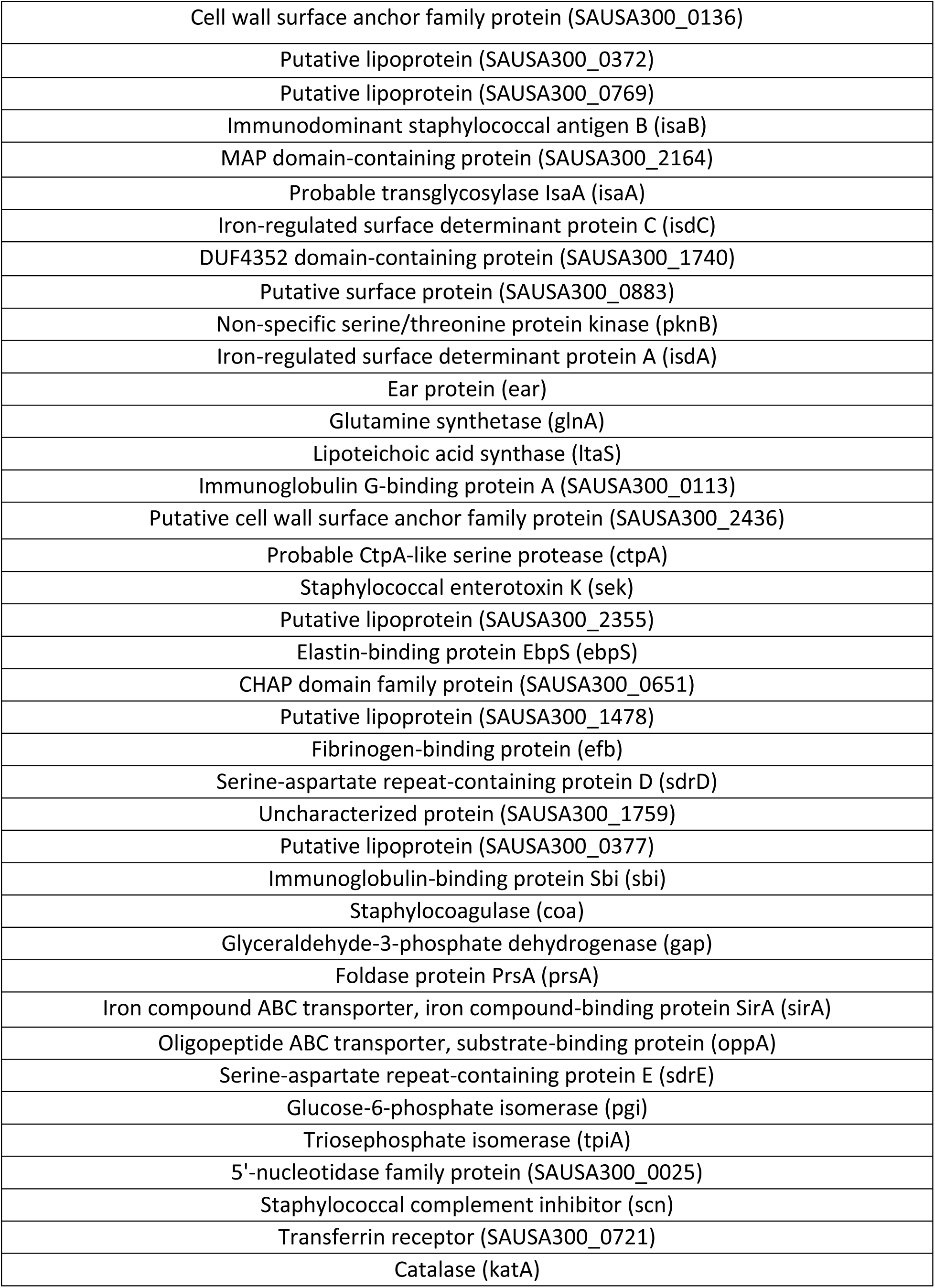
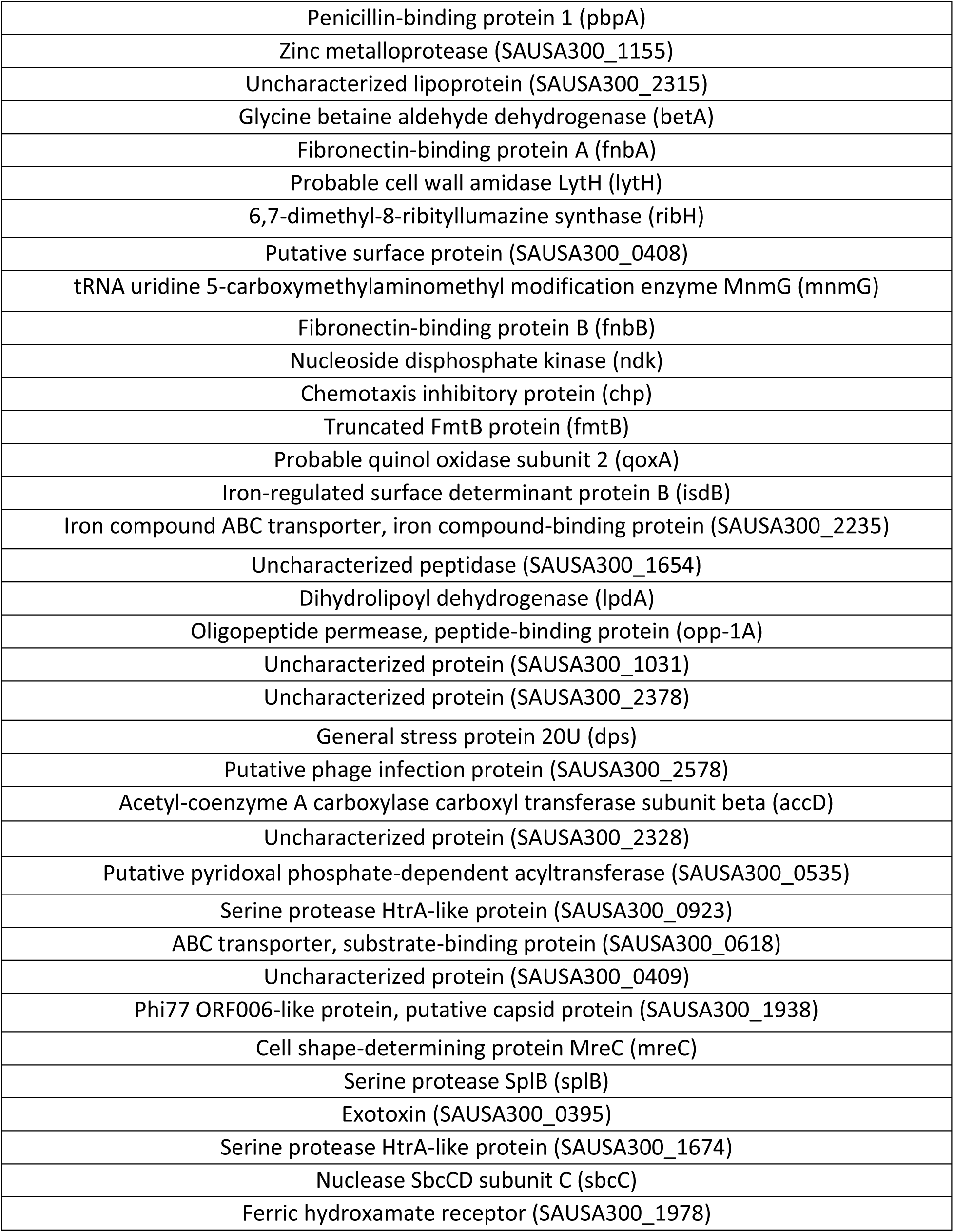
Mass spectrometry results of proteins with higher abundance in culture filtrate than cell pellets.

**Supplementary Table 2.**
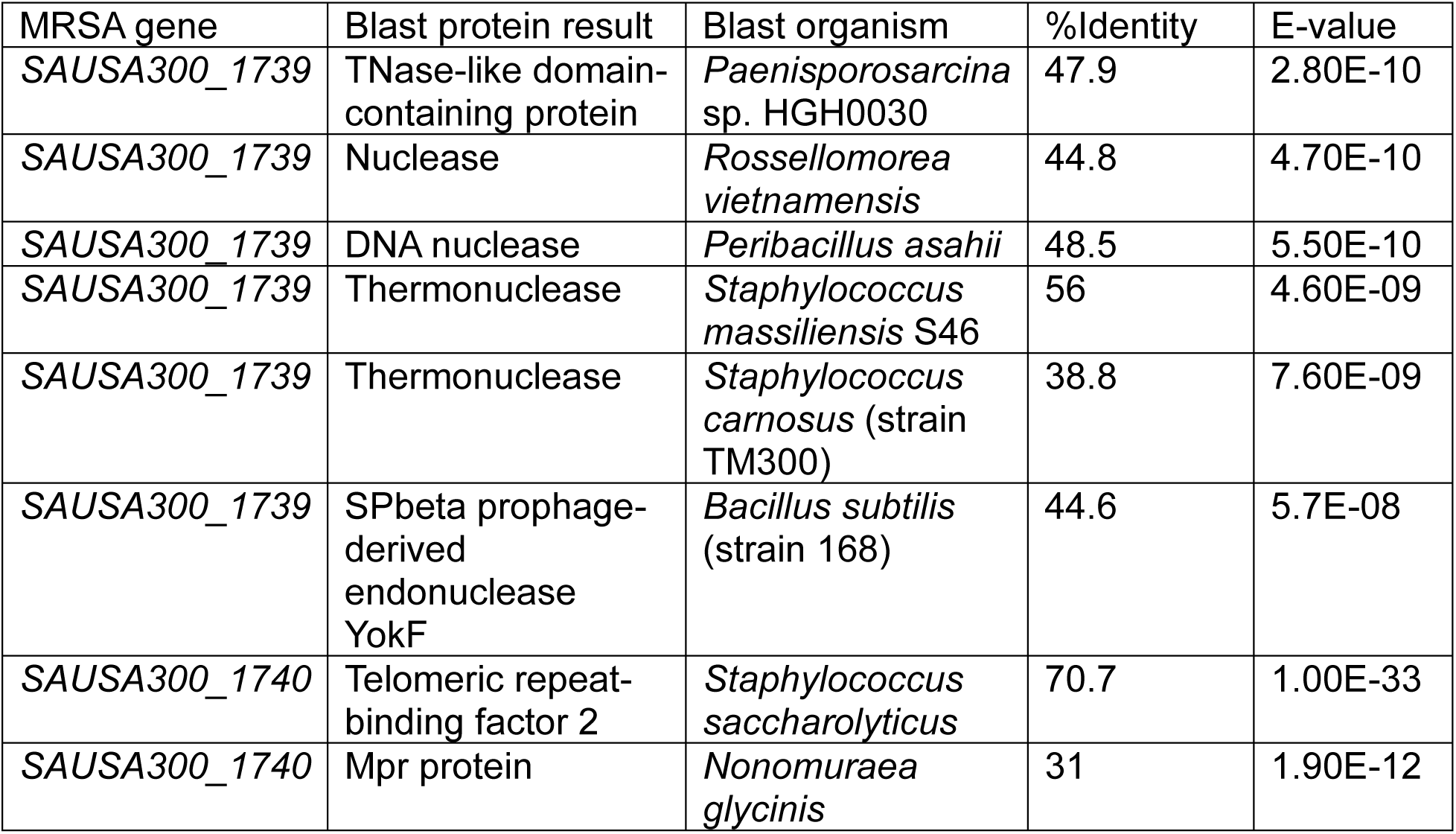
Select examples of protein blast homology results for mature lipoprotein sequences of *SAUSA300_1739* and *SAUSA300_1740*.

**Supplementary Figure 1.**
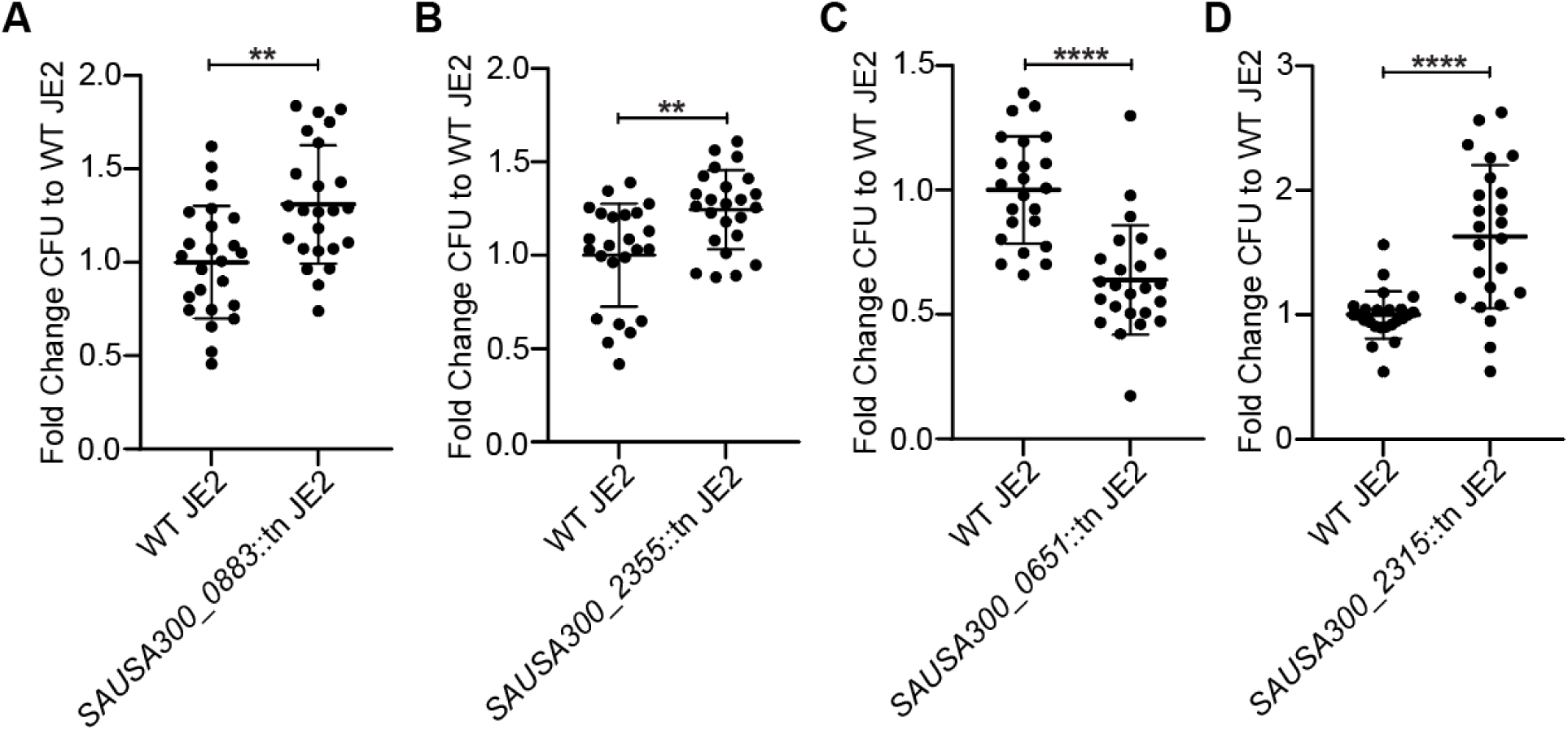
Transposon insertion mutants with differences in virulence compared to WT MRSA. MRSA USA300 JE2 were spinfected into WT C57BL/6 bone-marrow-derived macrophages (BMDMs) at a MOI of 10 followed by gentamicin protection for intracellular MRSA CFU, or BMDMs were infected without spinfection with MRSA at MOI of 10 for Lactate Dehydrogenase (LDH) release assay. **(A)** Intracellular CFU of WT and *SAUSA300_0883*::tn MRSA USA300 JE2 after 20 hours post spinfection. **(B)** Intracellular CFU of WT and *SAUSA300_2355*::tn MRSA USA300 JE2 after 20 hours post spinfection. **(C)** Intracellular CFU of WT and *SAUSA300_0651*::tn MRSA USA300 JE2 after 20 hours post spinfection. **(D)** Intracellular CFU of WT and *SAUSA300_2315*::tn MRSA USA300 JE2 after 20 hours post spinfection. Fold change CFU results in A-D are from comparison of individual biological replicates against the average WT MRSA USA300 JE2 CFU within the same experiment. Each transposon-inserted mutant results for A-D are from fold change results of 3 pooled experiments. **, p < 0.01; ****, p < 0.0001 (unpaired nonparametric Mann-Whitney U test).

**Supplementary Figure 2.**
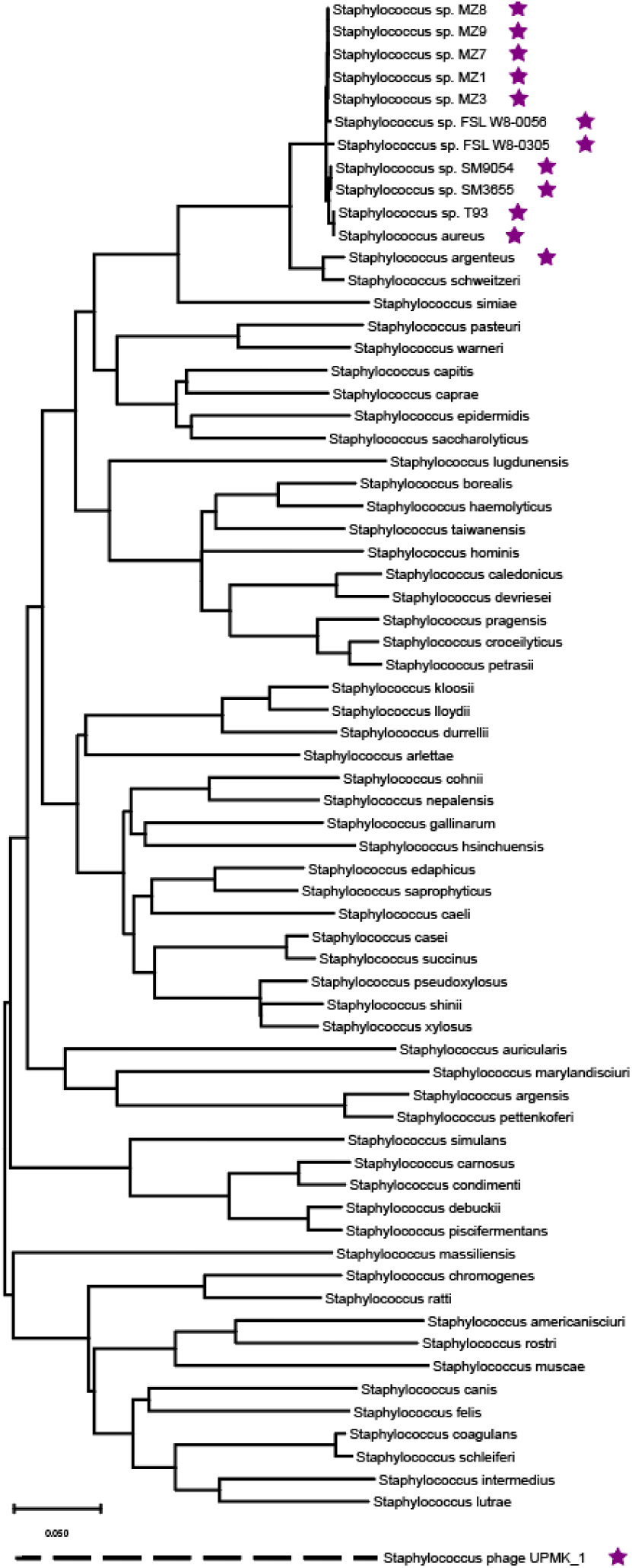
Genetic conservation of SAUSA300_1739 to SAUSA300_1744 operon in the *Staphylococcus* genus. Species phylogeny was constructed by extracting dnaA nucleotide coding sequence for each species. Sequences were aligned using MUSCLE and minimum evolution phylogeny was generated and tested with 50 bootstraps using MEGA 11. Species encoding *SAUSA300_1739* to *SAUSA300_1744* operon were identified using tBLASTn and confirmed by manual inspection of locus organization. Species and phage encoding the operon are indicated by presence of purple star next to name.

**Supplementary Figure 3.**
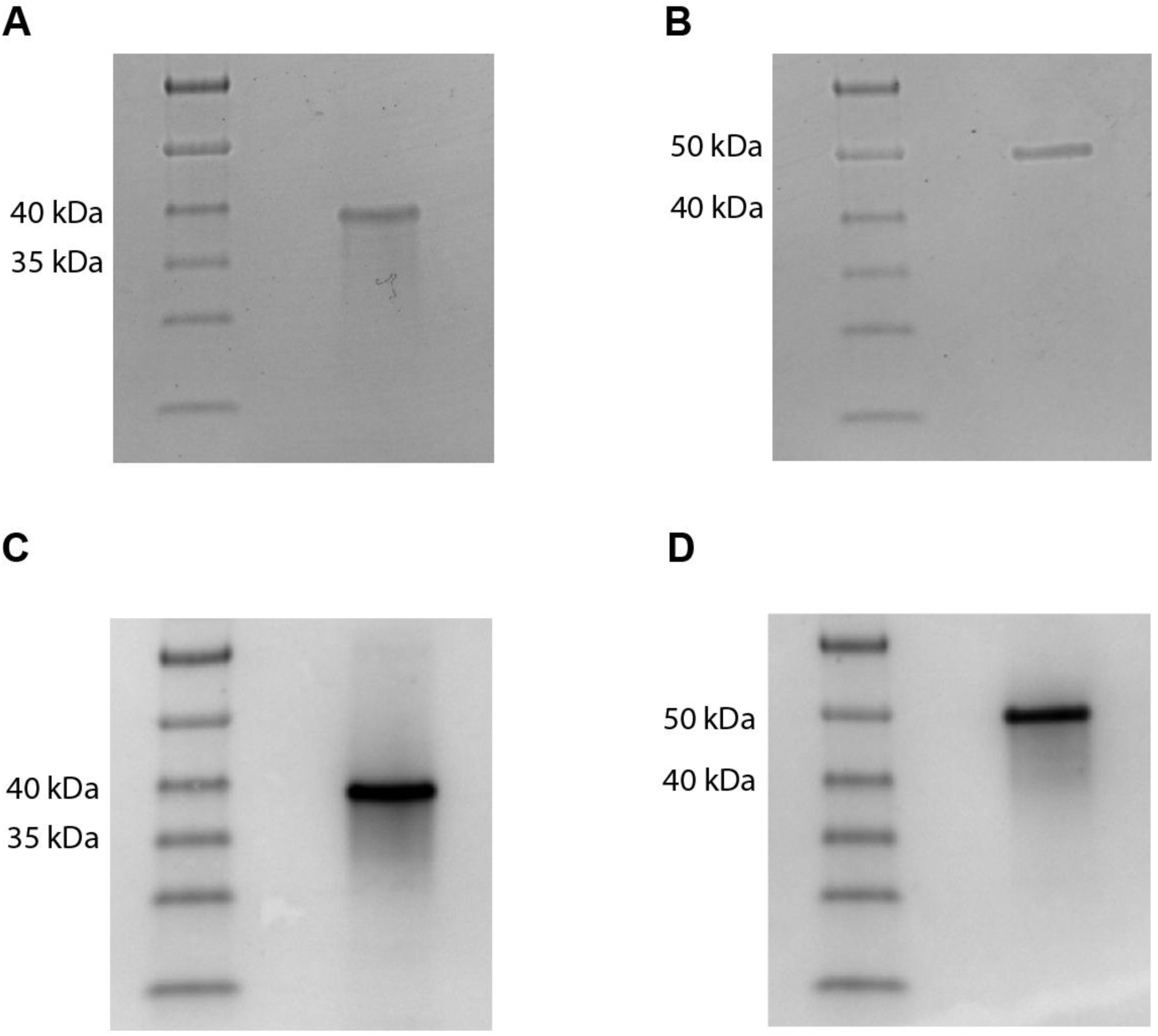
SDS-PAGE and western blot of purified SAUSA300_1739 and SAUSA300_1740. 1 μg of purified SAUSA300_1739-6xHIS-tag or SAUSA300_1740-6xHis-tag ran on SDS-PAGE gels and transferred onto PVDF membranes for western blot analysis with anti-His-tag antibody conjugated with HRP. **(A)** 1 μg of purified SAUSA300_1739-6xHIS-tag on SDS-PAGE gel with gel code blue stain. **(B)** 1 μg of purified SAUSA300_1740-6xHIS-tag on SDS-PAGE gel with gel code blue stain. **(C)** 1 μg of purified SAUSA300_1739-6xHIS-tag on PVDF membrane probed with anti-6xHis-tag antibody-conjugated HRP and stained with Opti-4CN. **(D)** 1 μg of purified SAUSA300_1740-6xHIS-tag on PVDF membrane probed with anti-6xHIS-tag antibody-conjugated HRP and stained with Opti-4CN.

**Supplementary Figure 4.**
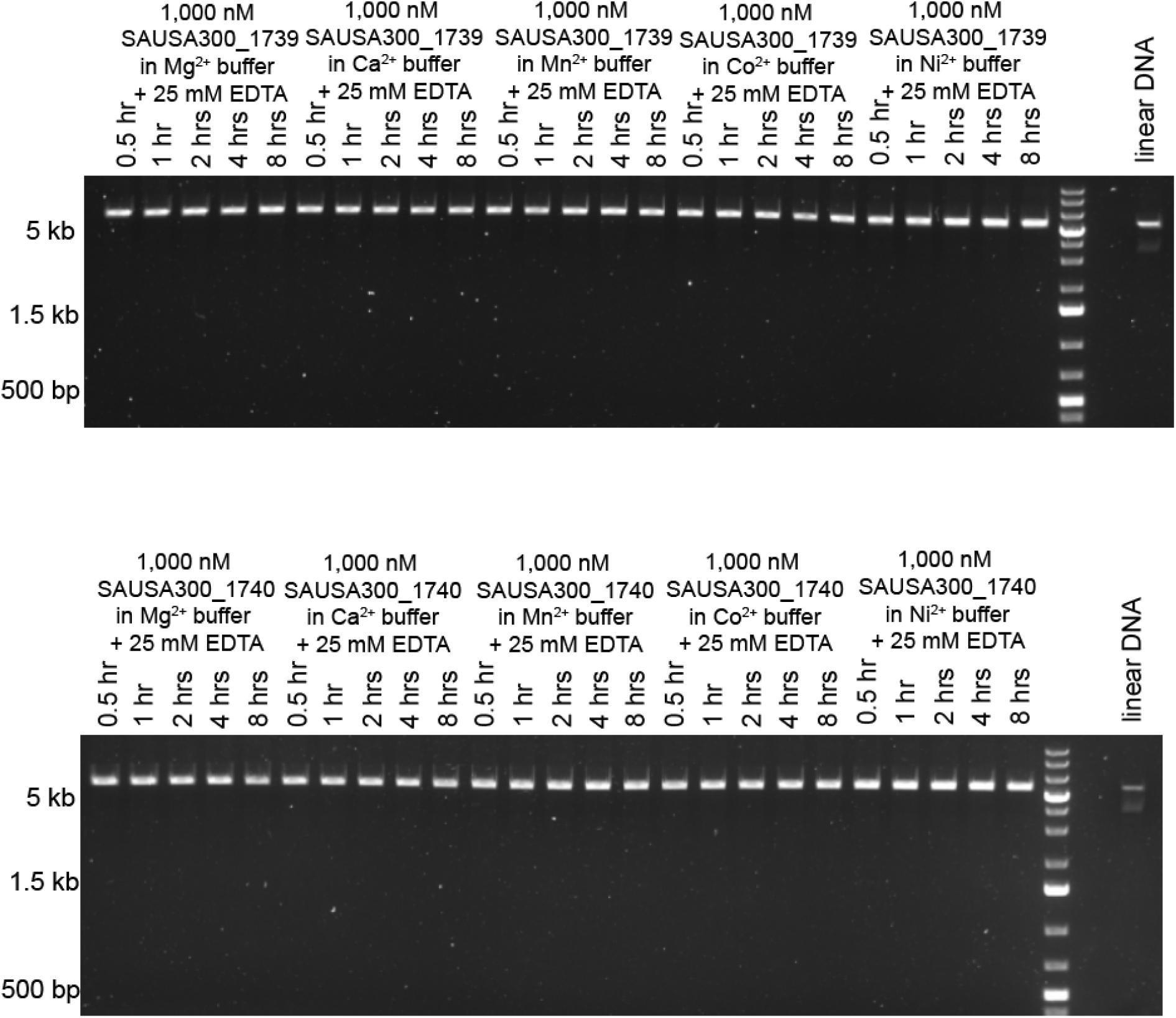
SAUSA300_1739 and SAUSA300_1740 have DNase activity. 1,000 nM purified unacylated SAUSA300_1739 or 1,000 nM purified unacylated SAUSA300_1740 was incubated with 100 ng of linear modified pRN11 plasmid (6 kb size) for 0.5, 1, 2, 4, or 8 hours at 37 °C in 3 mM Mg^2+^, Ca^2+^, Mn^2+^, Co^2+^, or Ni^2+^ containing buffer with 25 mM EDTA. Each agarose gel image is representative of 3 independent experiments.

